# Dynamics of Brain Connectivity across the Alzheimer’s Disease Spectrum: a magnetoencephalography study

**DOI:** 10.1101/2024.03.14.584574

**Authors:** Martín Carrasco-Gómez, Alejandra García-Colomo, Jesús Cabrera-Álvarez, Ricardo Bruña, Andrés Santos, Fernando Maestú

## Abstract

Alzheimer’s disease (AD) represents a major challenge in neurodegenerative disease research, as it is characterized by a complex pathophysiology that involves not only structural but also functional changes in the brain. While changes in static functional connectivity have already been linked to AD, there is still a lack of research studying dynamic functional connectivity (dFC) across the AD continuum, which could be crucial for identifying potential biomarkers for early diagnosis and tracking disease progression. This study leverages the high temporal resolution of MEG to dissect the dynamics of brain connectivity alterations across various stages of AD and their association with cognitive decline and structural brain changes.

321 participants were included in this study, categorized into healthy control, subjective cognitive decline (SCD), and mild cognitive impairment (MCI) groups. Amplitude envelope correlation with leakage correction was calculated over MEG signals using a sliding window, and the correlation across epochs was studied to assess dFC at whole-brain and node level. Finally, we explored dFC associations with cognitive scores, grey matter volume, and white matter fractal anisotropy.

The study unveils a significant reduction in whole-brain dFC, especially within the alpha and beta frequency bands, as individuals advance along the AD continuum. Notably, the frontal and temporal lobes, as well as regions within the default mode network, exhibited pronounced dFC reductions. Finally, dFC significantly correlated with cognitive performance and changes in structural brain, suggesting the potential of the proposed dFC metric as sensitive indicator for monitoring disease progression.

This investigation provides crucial insights into the temporal dynamics of brain connectivity alterations in the early stages of the AD spectrum, underlining the importance of dFC changes as reflective of cognitive and anatomical degeneration. The findings hint towards a strong relationship between connectivity profiles and white matter integrity, especially for high frequency activity in the association cortices.

**Key Points:**

- Dynamic functional connectivity declines over the AD spectrum.
- Dynamic functional connectivity reductions are most prominent in orbitofrontal, temporal, and DMN-related areas.
- Cognitive performance, brain volumetrics, and white matter integrity parameters positively correlate with dynamic functional connectivity over the whole AD spectrum, and most significantly among mild cognitive impairment participants.

## Introduction

Alzheimer’s disease (AD) is the most common form of dementia,^1^ affecting an estimated of 32 million people worldwide. In AD, functional networks supporting cognition become altered, including activation and deactivation deficits, abnormal oscillatory activity, and synaptic failure, and leading to impaired memory, behavioral disorders, and emotional changes.^2^

One way to evaluate the integrity of neural communication in the brain is through functional connectivity (FC), the statistical dependency of distant neural activity, which has been extensively used in the study of AD. More specifically, FC changes have been described along the disease spectrum, including alterations among individuals in the dementia phase of AD;^3^ in participants with mild cognitive impairment (MCI);^4,5^ in subjective cognitive decline (SCD),^6,7^ a preclinical asymptomatic stage in which participants already report a decrease in their cognitive capabilities, prodromal to MCI; and even as early as in healthy adults at risk of developing dementia.^8,9^ Taken together, these studies show an inverted u-shape pattern of FC changes along the continuum of AD, finding increased FC in the early stages, followed by a decrease in connectivity later on in the continuum.^10^ These changes have been previously associated with the underlying pathology hallmarks of AD.^11,12^

Notably, studies on brain connectivity have assumed, until recently, the stationarity of FC. However, the brain’s functional activity, far from being static, has been described as highly dynamic, forming various functional networks that underlie complex cognition.^13^ Consequently, by averaging over lengthy imaging recordings, transient information may be lost, and what we interpreted as inter-subject variability might in fact be averaged temporal variability. This perspective has brought attention to the field of spatiotemporal dynamics of brain networks, also called “dynamic functional connectivity” (dFC), which studies how FC evolves over time and what connectivity profiles arise. It has been proven that dFC changes dynamically in response to different contexts or stimuli, even in the resting-state.^14^

Given the consistency of the changes in static FC (sFC) throughout the continuum of AD, a novel line of research, focused on the possible dFC alterations underlying sFC changes, has become especially relevant for the disease. Most investigations involving dFC in AD have been carried out using functional magnetic resonance imaging (fMRI). A common finding in these studies is that dFC is reduced in AD, in key networks and regions for the disease, such as the DMN.^15^ Additionally, some studies have also reported such a decrease in dFC among MCI participants in regions like the precuneus, which correlated with cognitive performance.^16–18^ Likewise, Jones *et al*.^19^ found a reduction of time spent in connectivity states with a high contribution of posterior DMN areas among AD patients, and Schumacher *et al*.^20^ reported that AD patients spend more time in states of sparse connectivity, an inability to switch between states of low inter-network connectivity, and more highly and specifically connected network configurations than controls. For a review of fMRI findings about dFC in AD, see Filippi *et al*.^21^ and Matsui *et al*.^15^

However, the temporal resolution of fMRI is hampered by its low sampling frequency and the slow nature of the hemodynamic responses recorded, and is consequently likely to miss relevant dFC changes in the sub-second scale.^14^ Electrophysiological techniques such as electroencephalography (EEG) and magnetoencephalography (MEG) are able to overcome such difficulties, both because of their higher sampling frequency and the fact that they measure the electrical activity of cortical neurons. While very scarce, electrophysiological studies point in the same direction as those using fMRI, suggesting a decrease in dFC in the last stages of the AD continuum. Núñez *et al*.^14^ found decreased variance when studying the leaked-corrected amplitude envelope (AEC-c) correlations in sensor-space EEG in AD patients compared to the control group in both high alpha and low beta bands, while no significant differences appeared in the MCI group. In a posterior work they applied hidden Markov models to instantaneous FC in EEG sensor-space to study meta-states dynamics in AD and MCI, and found a decrease in modularity of meta-states in both MCI and AD participants compared to controls in both alpha and low beta bands,^22^ meaning that their resting-state networks were less well defined, possibly contributing to the shorter dwell times they also found in the pathological groups. In a functional near-infrared spectroscopy (fNIRS) and EEG integrative investigation, Li *et al*.^23^ found weaker or suppressed dFC in the high alpha and low beta bands of AD participants, reporting lower values for degree and clustering coefficient in the frontal pole and medial orbital regions of AD-induced brain networks. On the other hand, while not centered in dFC, other electrophysiological studies focused on other highly correlated parameters, such as entropy and dynamic changes in power, report findings in the same direction. Sun *et al*.^24^ compile, in a review, reports on decreasing complexity of the MEG/EEG signal in multiple entropy parameters in AD. Finally, Puttaert *et al*.^25^ applied hidden Markov models to MEG power topographies of healthy individuals and SCD, MCI, and AD participants, finding only differences in the dynamics of the control and AD groups. Similarly to the aforementioned fMRI study by Jones et al.,^19^ they found that AD participants spent less time in, and switched less frequently into, states with a high contribution of the posterior DMN.

To the best of our knowledge, no electrophysiological studies exist on the whole pre-dementia spectrum of AD that characterize the evolution of dFC in this population. Based on the presented premises, this study aims to characterize the evolution of dFC in the AD spectrum by estimating connectivity through AEC-c and correlating it over different segments of MEG data in a large cohort consisting of healthy controls, SCD participants, and both stable and progressive MCI participants. Guided by the claims of previous research, we hypothesize we will find the trend of dFC reduction earlier in the AD spectrum, as well as that these reductions to be predominant in frontal and temporal regions and in the DMN. Lastly, to wrap the interpretation of our findings, we will also study the correlation of dFC with cognitive scores, anatomical metrics, and white matter integrity markers, which we hypothesize to decline along with dFC.

## Materials and methods

### Participants

The sample for the present study consisted of 321 individuals with valid MEG resting state recordings, T1-weighted magnetic resonance imaging (MRI) scans, and a thorough neuropsychological evaluation. Of them, 143 were unimpaired older adults with no cognitive complaints, namely the healthy control (HC) group, 72 were unimpaired individuals with SCD, and 106 presented MCI.

MCI patients were diagnosed according to the National Institute on Aging-Alzheimer Association (NIA-AA) criteria,^26^ based on their cognitive profile. Besides meeting the core clinical criteria for MCI, patients also exhibited significant hippocampal atrophy according to the evaluation of an experienced neuroradiologist who was blinded to the clinical outcome (see also below hippocampal volumes calculation). Consequently, patients were categorized as “MCI due to AD with intermediate likelihood”.^26^ MCI participants underwent longitudinal assessment on their clinical progression after a mean of 2 (standard deviation of 0.5 years). They were further subdivided into 64 stable MCI (sMCI) and 42 progressive MCI (pMCI) patients, depending on their progression to probable AD in during the follow-up period.^27^ The participants were recruited from different Spanish institutions and associations; namely, the “Fulbright Alumni Association”, the “Asociación Española de Ingenieros de Telecomunicación”, or the “Hospital Universitario Clínico San Carlos”, being the last one located in Madrid.

To correctly characterize the SCD group, cognitive concerns were self-reported by the participants in an interview with clinician experts. The final group assignment was made after neuropsychological evaluation attending to a multidisciplinary consensus (by neuropsychologists, psychiatrists, and neurologists). To prevent possible confounders of SCD, problematic medication, psycho-affective disorders or other relevant medical conditions lead to the exclusion from the study. Following the recommendations made by the SCD-I-WG, all subjects were older than 60 at onset of SCD, and this onset took place within the last 5 years prior to the recruitment.

The exclusion criteria employed in this study were the followings: (1) history of psychiatric or neurological disorders or drug consumption that could affect MEG activity, such as cholinesterase inhibitors; (2) evidence of infection, infarction or focal lesions in a T2-weighted scan within 2 months before MEG acquisition; (3) history of alcoholism, chronic use of anxiolytics, neuroleptics, narcotics, anticonvulsants or sedative hypnotics; and (4) for the healthy individuals (HC and SCD groups), a score below 26 points in the MMSE. Besides, additional analyses were conducted to rule out other possible causes of cognitive decline, such as B12 vitamin deficit, diabetes mellitus, thyroid problems, syphilis, or infection by the human immunodeficiency virus (HIV).

All participants were native Spanish speakers and provided written informed consent. The Institutional Review Board Ethics Committee at Hospital Universitario San Carlos approved the study protocol, and the procedure was performed following the Helsinki Declaration and National and European Union regulations. Table 1 summarizes relevant demographic and clinical information of the sample, and Table S1 compares demographics from sMCI and pMCI groups.

**Table 1.**
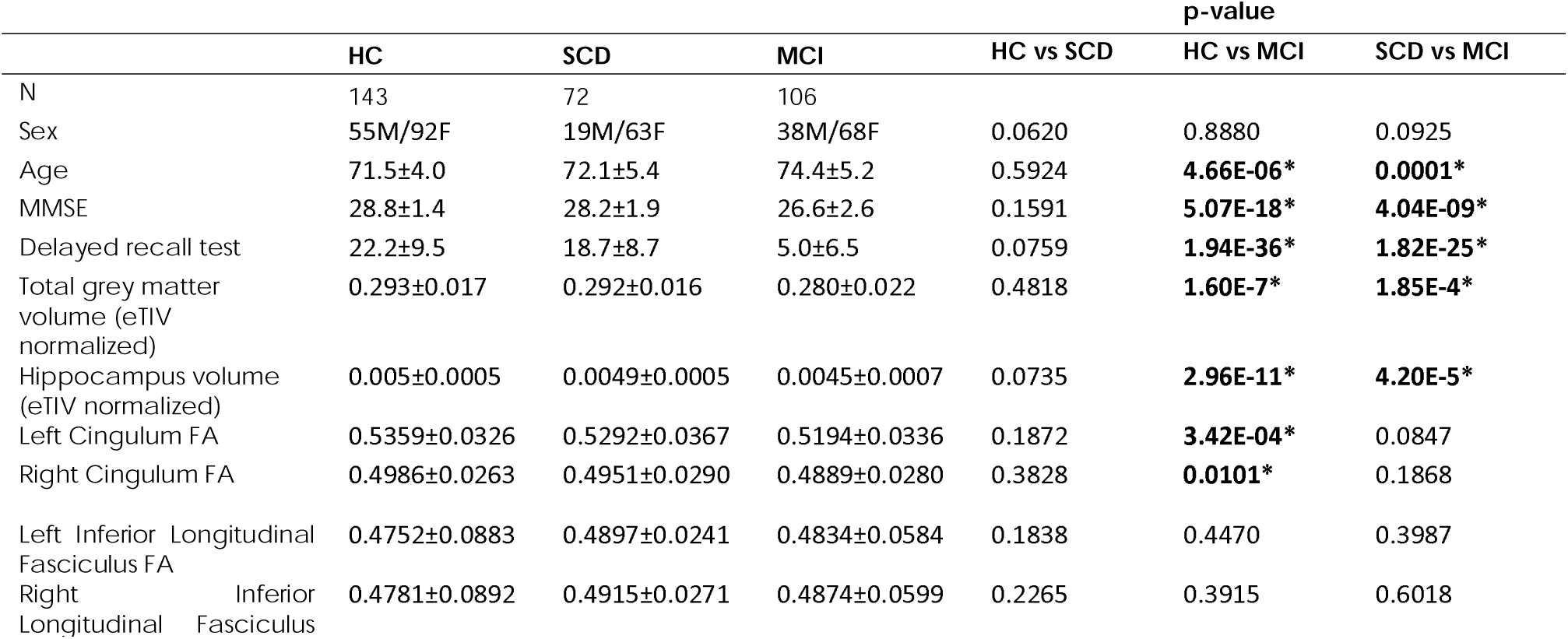
Sample demographics, neuropsychological test scores, and anatomic and white matter parameters. Pair-wise group-level differences are shown in the right columns, with significant results after FDR correction (q = 0.05) with an asterisk. HC: healthy control; SCD: subjective cognitive decline; MCI: mild cognitive impairment; MMSE: Mini Mental State Examination; FA: fractional anisotropy.

### Neuropsychological assessment

To assess the general cognitive and functional status of the participants, the following set of screening questionnaires were used: The Mini Mental State Examination (MMSE);^28^ the Geriatric Depression Scale–Short Form (GDS);^29^ the Hachinski Ischemic Score, (HIS);^30^ and the Functional Assessment Questionnaire (FAQ).^31^ After the initial screening, all subjects underwent an exhaustive neuropsychological assessment including: Direct and Inverse Digit Span tests (Wechsler Memory Scale, WMS-IV);^32^ Immediate and Delayed logic memory component of the WMS-IV;^32^ and Phonemic and Semantic Fluency (Controlled oral Word Association Test, COWAT).^33^ From these measures, both MMSE scores and the Delayed logic memory component of the WMS-IV were used in this study.

### Magnetic resonance imaging

An individual T1-weighted MRI scan was acquired for each participant in a General Electric 1.5 T scan. A high-resolution antenna was employed, together with a homogenization phased array uniformity enhancement filter (fast spoiled gradient echo sequence, TR/TE/TI of 11.2/4.2/450 ms; flip angle of 12°; slice thickness of 1 mm, 256 × 256 matrix, and FOV of 25 cm. These images were processed using the FreeSurfer software (version 6.1.0) for automated cortical and subcortical segmentation and parcellation.^34^

Additionally, diffusion weighted imaging (DWI) scans were acquired using a single shot echo planar imaging sequence with a TE of 96.1 ms, a TR of 12ms; a NEX 3 for increasing the signal-to-noise ratio, a FOV of 30.7 cm (resolution of 128 x 128), and a 4 mm slice thickness. In the DWI sequence, one image had no diffusion sensitization (i.e., T2-weighted b0 images), and 25 were DWI (b = 900 s/mm2). DWI images were processed with AutoPtx for probabilistic tractography, as outlined in the paper of de Groot *et al.*^35^ The procedure extracted the mean fractional anisotropy (FA) for each of the tracts of interest (see next paragraph).

The volumetric and tractography measures that were included in further analyses were the total volumes (mm^3^) of cortical gray matter and hippocampus, normalized by the estimated intracranial volume (eTIV), and the FA of the cingulate gyrus part of the cingulum (CGC) and the inferior longitudinal fasciculus (ILF), due to their relevance in the early progression of the disease.^36^

### MEG data acquisition

Each participant underwent a four-minute, eyes-closed, resting-state MEG scan at the Center for Biomedical Technology in Madrid, Spain. This session used a 306-sensor Elekta Neuromag system (Elekta AB, Stockholm, Sweden) to record brain activity. Additionally, ocular and cardiac activities were monitored using two sets of bipolar electrodes. To track head position, four head position indication coils were placed on the participants’ scalp: two on the forehead and two on the mastoids. These coils, along with approximately 200 points of the participant’s head shape, were digitized using the Fastrak 3D scan (Polhemus, Colchester, VT, USA). The coils were actively used during the recording to continuously monitor the head’s position in relation to the MEG helmet. The procedure was conducted in a magnetically shielded room, and participants were advised to remain as motionless and relaxed as possible. The data were filtered in real time between 0.1 and 330 Hz and digitized at a rate of 1000 Hz. For data processing, the spatiotemporal signal space separation method,^37^ as implemented in the Neuromag software (MaxFilter version 2.2, with a correlation of 0.90 and a time window of 10 seconds), was employed to eliminate external noise and correct for any head movements during the MEG scan.

### MEG preprocessing and source reconstruction

MEG data were blindly preprocessed by an electrophysiology expert using FieldTrip software.^38^ This process involved dividing the continuous data into non-overlapping, artifact-free 4-second segments, with an extra 2 seconds of real data to each side for padding purposes. Participants who had at least 20 valid segments were selected for further analysis. Due to the high redundancy in the data after applying the signal space separation method,^39^ analysis focused solely on data from the magnetometers.

Each participant’s T1-weighted MRI scan was used for individual source reconstruction. The source model was established in MNI space using a uniform 3D grid with 10 mm spacing. Source positions were identified based on the automated anatomical labeling atlas,^40^ resulting in 1210 cortical sources distributed across 80 regions of interest. This source model was then linearly transformed to each participant’s T1-weighted scan. The scans were segmented using SPM12 software,^41^ creating a brain mask. This mask aided in constructing the participants’ single-shell realistic head models. These models, combined with the participants’ source models, facilitated the generation of a lead field using a modified spherical solution.^42^

To derive time series data for each source location, a linearly constrained, minimum variance beamformer was used as the inverse model. To this end, the beamformers were computed using the covariance matrix of the data filtered between 2 and 45 Hz, with a regularization level set at 1% of the average sensor power.

### Dynamic functional connectivity

#### dFC calculation

FC was estimated using the AEC with leakage correction,^43^ either between each pair of time-series in the 80 ROIS of the AAL or between the 1210 brain sources, using the absolute value of the Pearson correlation between the envelopes of the band-pass filtered time-series. Data was filtered in the theta (4-8 Hz), alpha (8-12 Hz), and beta (12-30 Hz) frequency bands through a finite impulse response filter with an order of 1800, designed with a hamming window. AEC was chosen because of its reliability, its within and between subject consistency,^44^ and due to its sensibility to dFC in aforementioned literature with a similar objective.^14,22,45^

Typically, this connectivity metric is calculated in each of the defined signal segments, and then averaged over those, obtaining the so-called static FC (Figure 1A). However, for the sake of studying dFC, we calculated the correlation between the connectivity matrices corresponding to each segment. This is usually labeled in literature as the recurrence matrix (C),^22,46^ and represents the similarity of connectivity distributions over time, as represented in figure 1B. To maintain a uniform representation of the data of all subjects, the same number of segments were used for dFC calculations in all subjects. Given that the minimum number of segments among the participants was 21, we just kept the first 21 clean segments of each participant.

**Figure 1.**
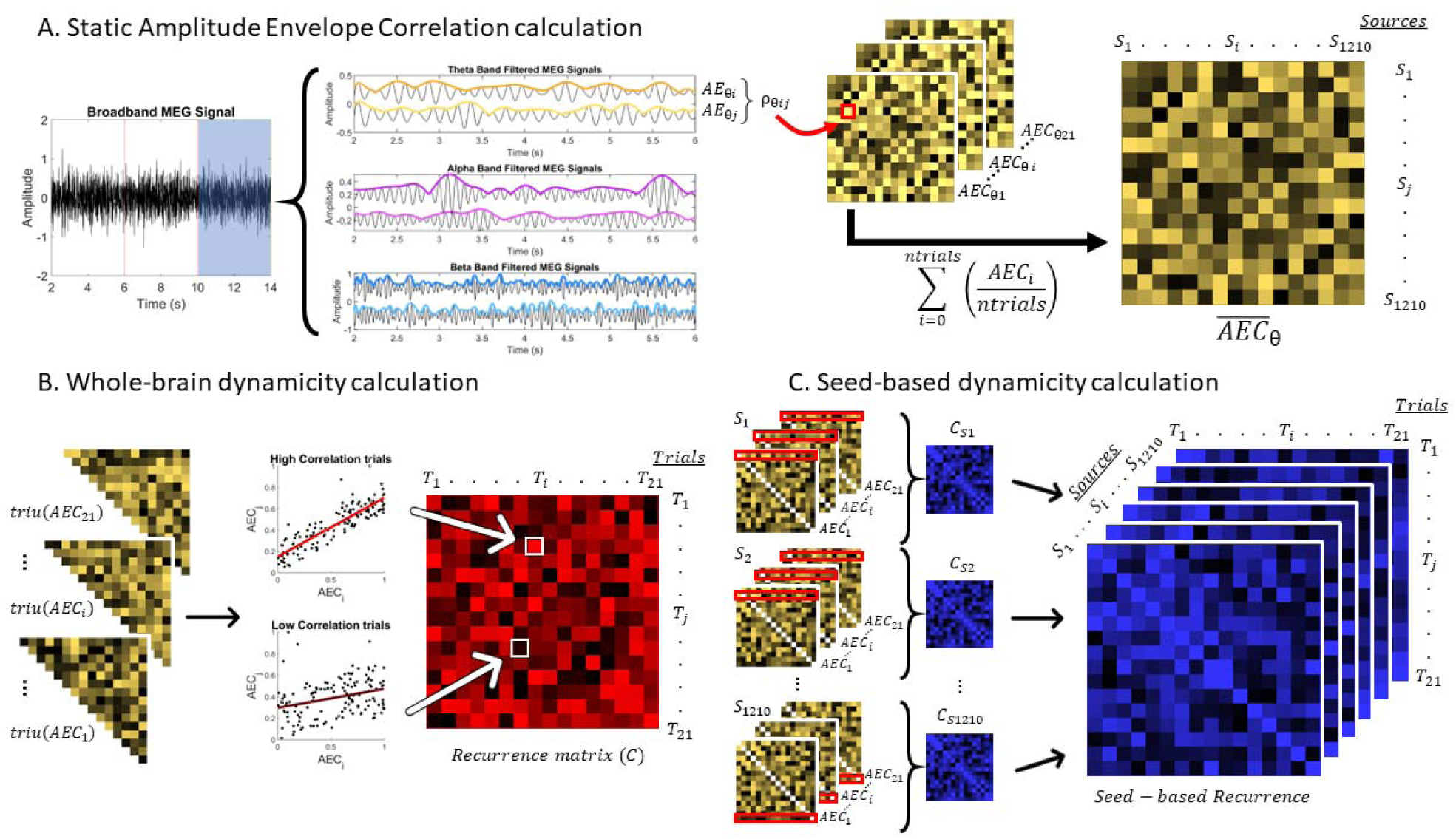
Depiction of the methods used in this study. (**A**) Description of the procedure to calculate AEC, including the signal filtering, envelope extraction through the Hilbert transform, correlation between envelopes, and averaging over trials to obtain the static functional connectivity measure of AEC. (**B**) Whole brain dynamicity calculation. Each of the AEC upper triangular matrices extracted from each signal segment of four seconds is then correlated with each other, obtaining a “number of trials” by “number of trials” recurrence matrices. The upper triangle of this matrix is then averaged to obtain a final dynamicity parameter for each subject. (**C**) Seed-based dynamicity calculation. In this case, and for each source, the connectivity of that source with the rest is extracted to obtain the recurrence matrices, obtaining a seed-based recurrence matrix of dimensions number of sources x number of trials x number of trials. Later, the upper triangle of each source recurrence matrix is averaged to get a dynamicity parameter for each of the sources, which enables us to extract spatial comparisons between groups.

#### Whole-brain dFC

Each of the C_n_ terms of the recurrence matrix represents the correlation of all connectivity terms of the connectivity matrix of segment *i* with those of the connectivity matrix of segment *j*, resulting in a matrix of dimensions number of segments × number of segments. Consequently, we obtained 321 21 × 21 recurrence matrices, one for each subject, representing the consistency of connectivity profiles over time. The Fisher Z-transformation was subsequently applied to these matrices, to prevent any possible skewness in the calculated distribution.^47^ In order to acquire a parameter representing the dynamic nature of the connectivity, we calculate dynamicity as:

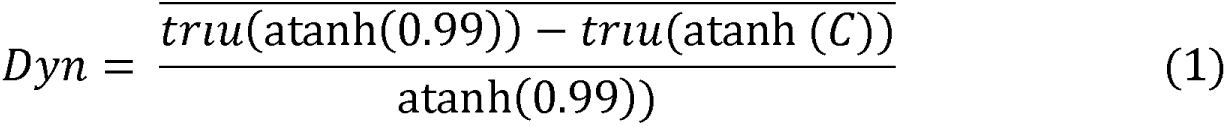

Where the operator *triu* is the upper triangle selector, the operator ’:f is the average of ’:f, the operator *atanh* is the inverse hyperbolic tangent, equivalent to the Fisher z-transform, and *C* is the recurrence matrix. This parameter would range from 0 to 1, where 0 represents complete stationarity, while 1 would represent a complete lack of temporal correlation among connectivity matrices. Note that we use the *atanh* of 0.99 instead of 1 for normalization, since the inverse hyperbolic tangent of 1 is infinite.

#### Spatial dFC analysis

No spatial information can be derived from the previous recurrence matrix. To perform a data-driven analysis of the spatial distribution of changes in dFC through recurrence matrices, we propose the methodology we called “seed-based recurrence matrix”. While all sources are used to calculate the correlation of each segment with the others in a recurrence matrix, the seed-based recurrence matrix separates that process for each of the sources, as shown in Figure 1C.

For each source, the connectivity values of that source with the remaining sources are extracted, and then correlated along the different segments, obtaining a number of segments × number of segments recurrence matrix for each of the original 1210 source positions. Consequently, the seed-based recurrence matrix has a dimension of sources × number of segments × number of segments. Once again, the Fisher Z-transformation was applied to the data to favor the normality of the distribution and the result was subtracted from the inverse hyperbolic tangent of 0.99, and normalized by the same number, as in Equation 1. Given the number of sources used for the source reconstruction, we obtained 321 1210 × 21 × 21 seed-based recurrence matrices, one for each participant. Equally as before, we averaged the upper triangle values of the recurrence matrices of each source to finally obtain 1210 values per subject.

### Statistical analyses

Initially, demographic characteristics were evaluated, using an independent samples t-test to compare age and neuropsychological variables, as well as a chi-square test to compare sex proportion between each of the groups.

#### Whole-brain dFC

Statistical analyses were carried out in a stepwise manner. Firstly, we studied whole-brain dFC across the different groups in our sample, to study whether global dFC diminishes along the AD continuum. To this end, we firstly compared the dynamicity values (Equation 1), between groups in each frequency band (theta, alpha and beta) through a regression analysis, introducing sex and age as covariates. FDR correction was applied to each frequency band set of comparisons through the Benjamini-Hochberg procedure,^48^ with a *q* of 0.05. Any *p-values* surviving FDR were considered statistically significant, and β*-values* were used for interpreting the directionality of results.

Given that we also had the information on the progression towards AD of the participants in the MCI group, we also carried out an additional test comparing the whole-brain dFC of the participants in the sMCI and pMCI groups.

#### Spatial dFC analysis

The second step of our analysis aimed to study the spatial distribution of dFC changes between groups. For that purpose, the aforementioned seed-based recurrence matrices (Figure 1C) of each subject were used. To account for multiple comparisons, we conducted two-sided nonparametric cluster-based permutation tests, using a Montecarlo approach and an independent samples F-statistic ANOVA in each sample, including sex and age as covariates. The source-level and cluster-level significance thresholds were set to 0.05. Significant sources were grouped based on spatial contiguity, obtaining a cluster statistic defined as the sum of the individual statistics of each source in it. We conducted 10,000 permutations on the original data to create a surrogate distribution of random cluster statistics to compare with our original result. As this test was a pos-hoc analysis, it was only performed between groups that showed significant differences in global dFC (i.e., in the previous analysis). Subsequently, a post-hoc linear regression analysis comparing the seed-based dynamicity between groups at the found cluster was performed to check for the directionality of the results.

#### Neuropsychological, volumetric and DTI FA correlations

Finally, we investigated whether the observed changes in dFC were related to changes in cognition, brain volumetrics, and white matter integrity. To do so, we calculated Pearson’s partial correlation of the dynamicity with the previously mentioned neuropsychological test scores, volumetry parameters, and tract FA in the frequency bands where statistical differences in global dFC were detected. We performed these tests both using the data from all groups as well as only from the individual groups, including sex and age as covariate in all cases. FDR correction was again applied to each frequency band set of comparisons, with a *q* of 0.05.

## Results

### Sample demographics and neuropsychological assessment

Sample demographics and neuropsychological test scores, as well as group-level differences, are shown in Table 1, and in Supplementary Table 1 for sMCI and pMCI groups.

The age of the MCI group was significantly higher than that of both SCD and HC groups, and while sex did not significantly differ between any of the groups, *p*-values were close to significance, supporting the inclusion of both variables as covariates in the rest of statistical tests of the study. MCI performed significantly worse than both HC and SCD groups in all the neuropsychological tests.

### Whole-brain dFC is reduced along the AD spectrum

Whole-brain dFC was compared through dynamicity (Equation 1) using a linear regression test between each pair of groups, including sex and age as covariates, as shown in Figure 2. Dynamicity values in the HC group were significantly higher than those in the MCI group after FDR correction (*q* = 0.05) in both alpha (*corrected p-value* = 0.0002, β*-value* = -0.0265, *Cohen’s d* = 0.474) and beta (*corrected p-value* = 0.0008, β*-value* = -0.0135, Cohen’s d = 0.506). On the other hand, the SCD group lies in the middle, with nonsignificant differences with neither the HC nor the MCI group (all *corrected p-values* and β*-values* can be found in Supplementary Table 2). Post-hoc comparisons between sMCI and pMCI groups were all non-significant, as shown in Supplementary Figure 1 and Supplementary Table 3.

**Figure 2.**
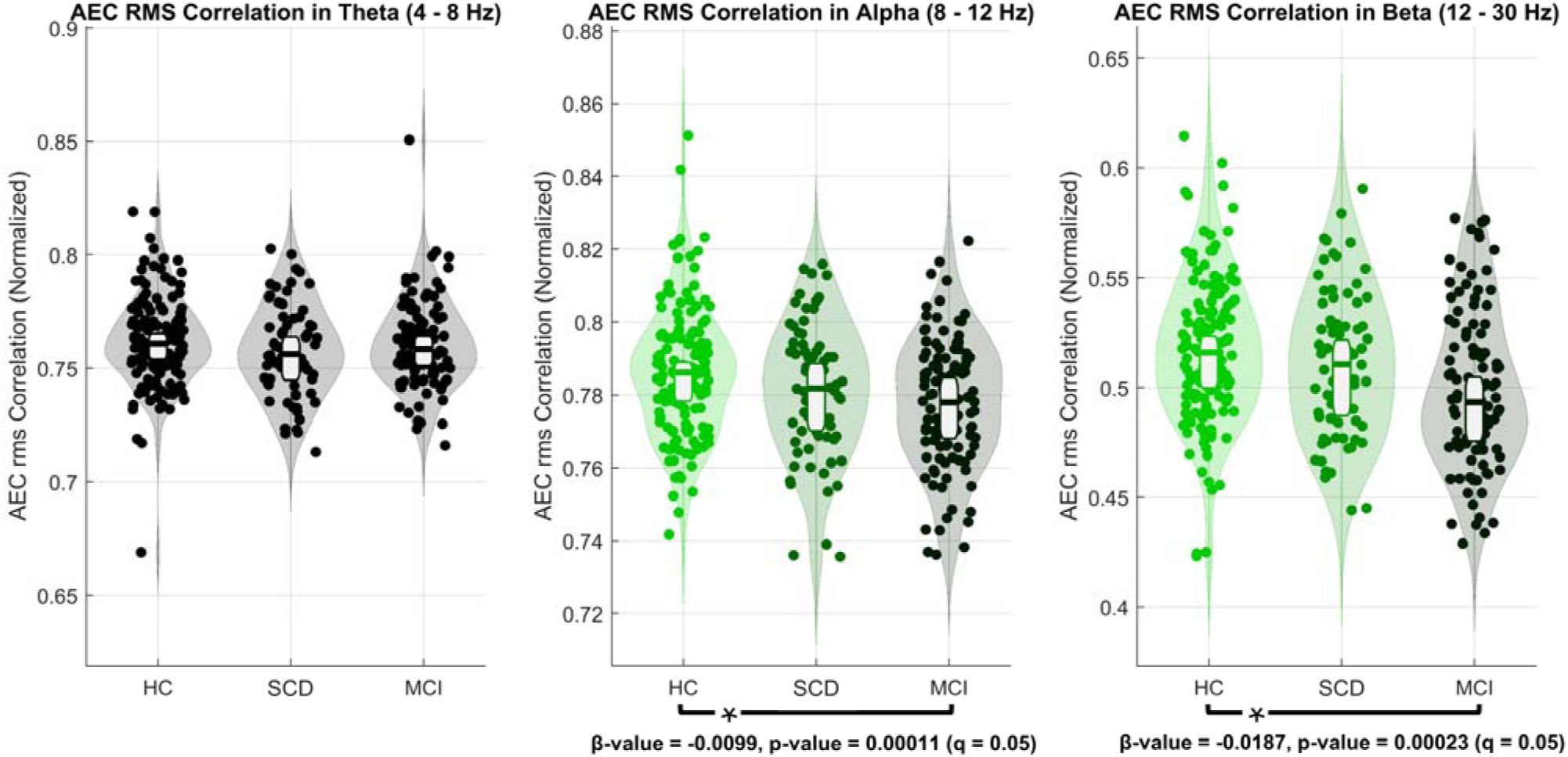
Whole-brain dynamicity comparisons between healthy control (HC), subjective cognitive decline (SCD) and mild cognitive impairment (MCI) groups. Significant differences were seen between the HC and MCI groups in the alpha (*corrected p-value* = 0.00015, β*-value* = -0.0265, *Cohen’s d* = 0.474, q = 0.05) and beta (*corrected p-value* = 0.00076, β*-value* = -0.0135, *Cohen’s d* = 0.506, q = 0.05). No differences were observed in the theta band or between the SCD group and the rest in any frequency band.

### Localization of dFC changes along the AD continuum

Seed-based dynamicity was compared through nonparametric CBPTs between those groups with significant differences in whole-brain dFC, namely between the HC and MCI groups in the alpha and beta frequency bands. Significant statistical differences were found in seed-based dynamicity between the HC and MCI groups in both the alpha (*p-value* = 0.0288) and the beta (*p-value* = 0.009) bands.

The cluster showing the most notable differences in the alpha band (Figure 3A) included the orbital and medial-orbital areas of the frontal gyrus, the left hippocampus and parahippocampus, as well as the left temporal pole and parts of the inferior and middle temporal gyrus. A post-hoc linear regression analysis found that MCI participants showed a significant reduction in seed-based dynamicity compared to HC in this cluster (Figure 3B; *p-value* < 0.0001; β*-value* = -0.0506; *Cohen’s d* = 0.594). For a complete list of the AAL ROIs contained in the cluster, please see Supplementary Table 4.

**Figure 3.**
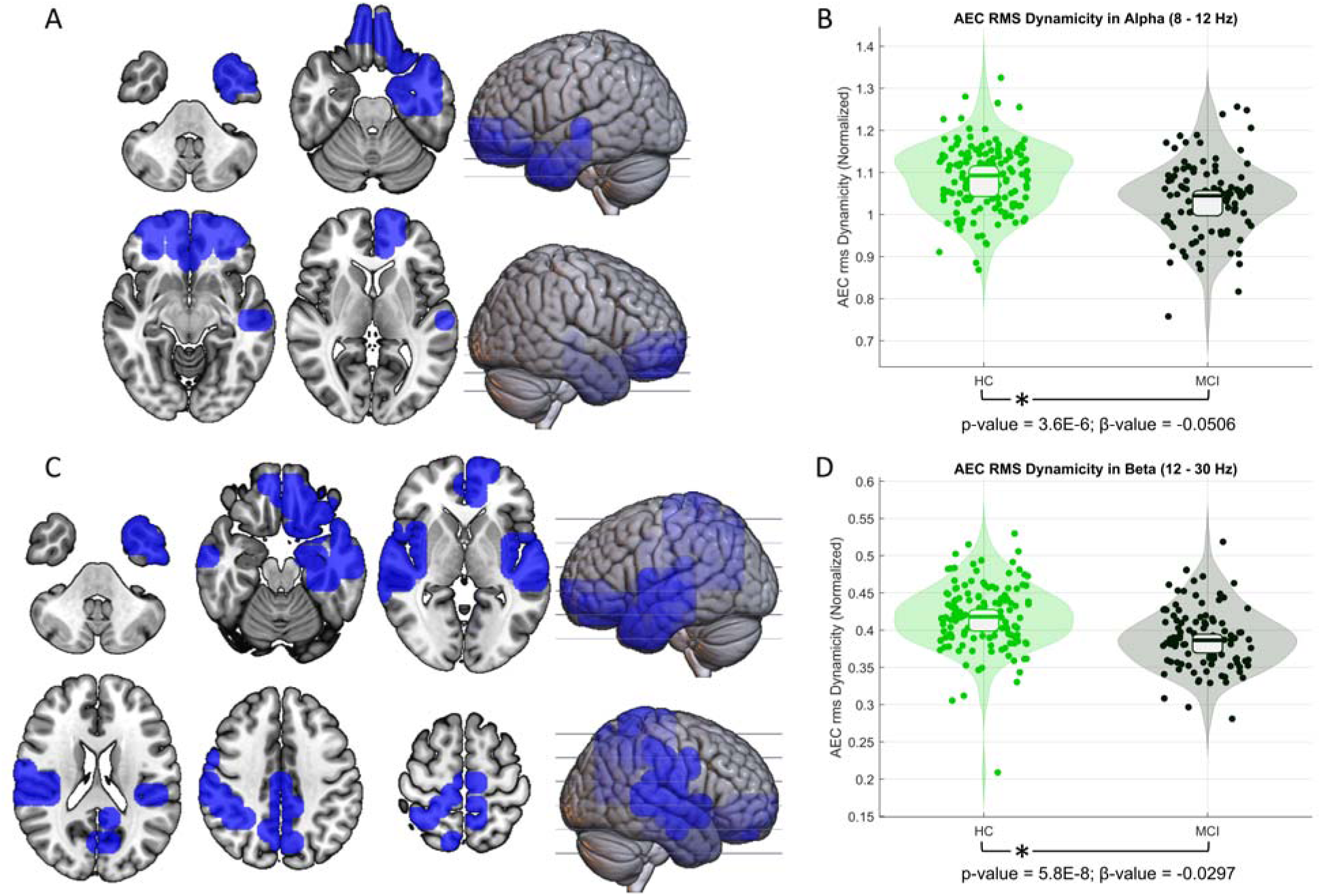
Results of the CBPT analyses for seed-based dynamicity. These comparisons were performed between those groups with significant differences in whole-brain dynamicity. Thus, CBPT tests were performed between the MCI and HC groups in the alpha and beta bands. (**A**) Significant cluster showing altered dynamic functional connectivity (dFC) in the alpha band (*p-value* = 0.0285). (**B**) Linear regression test results for the alpha band seed-based dynamicity values in the significant cluster depicted in A, showing a significant decrease of dFC in the MCI group (*Cohen’s d* = 0.594). (**C**) Significant cluster showing altered dFC in the beta band (*p-value* = 0.0089). (**D**) Linear regression test results for the alpha band seed-based dynamicity values in the significant cluster depicted in C, showing a significant decrease of dFC in the MCI group (*Cohen’s d* = 0.668).

In the beta band, the cluster showing maximal differences (Figure 3C) comprised the left orbital and right superior frontal gyrus, left temporal pole, bilateral temporal gyrus, left hippocampus and parahippocampus, the anterior, middle, and posterior cingulate gyrus, the precuneus, the left inferior parietal gyrus, and some parts of the calcarine fissure, fusiform gyrus, and cuneus. Again, the post-hoc linear regression analysis found a significant reduction in the MCI group compared to HC in the mentioned cluster (Figure 3D; *p-value* < 0.0001; β*-value* = -0.0297; *Cohen’s d* = 0.668). For a complete list of the AAL ROIs contained in the cluster, please see Supplementary Table 5.

### dFC correlates with cognitive performance along the AD continuum and in MCI

Finally, we investigated the correlation between dFC and cognition, brain volume and DTI parameters, both considering the whole dataset and the individual groups. Figure 4 depicts correlations between dynamicity and MMSE, eTIV-normalized hippocampal volume, and fractal anisotropy of the left cingulum.

**Figure 4.**
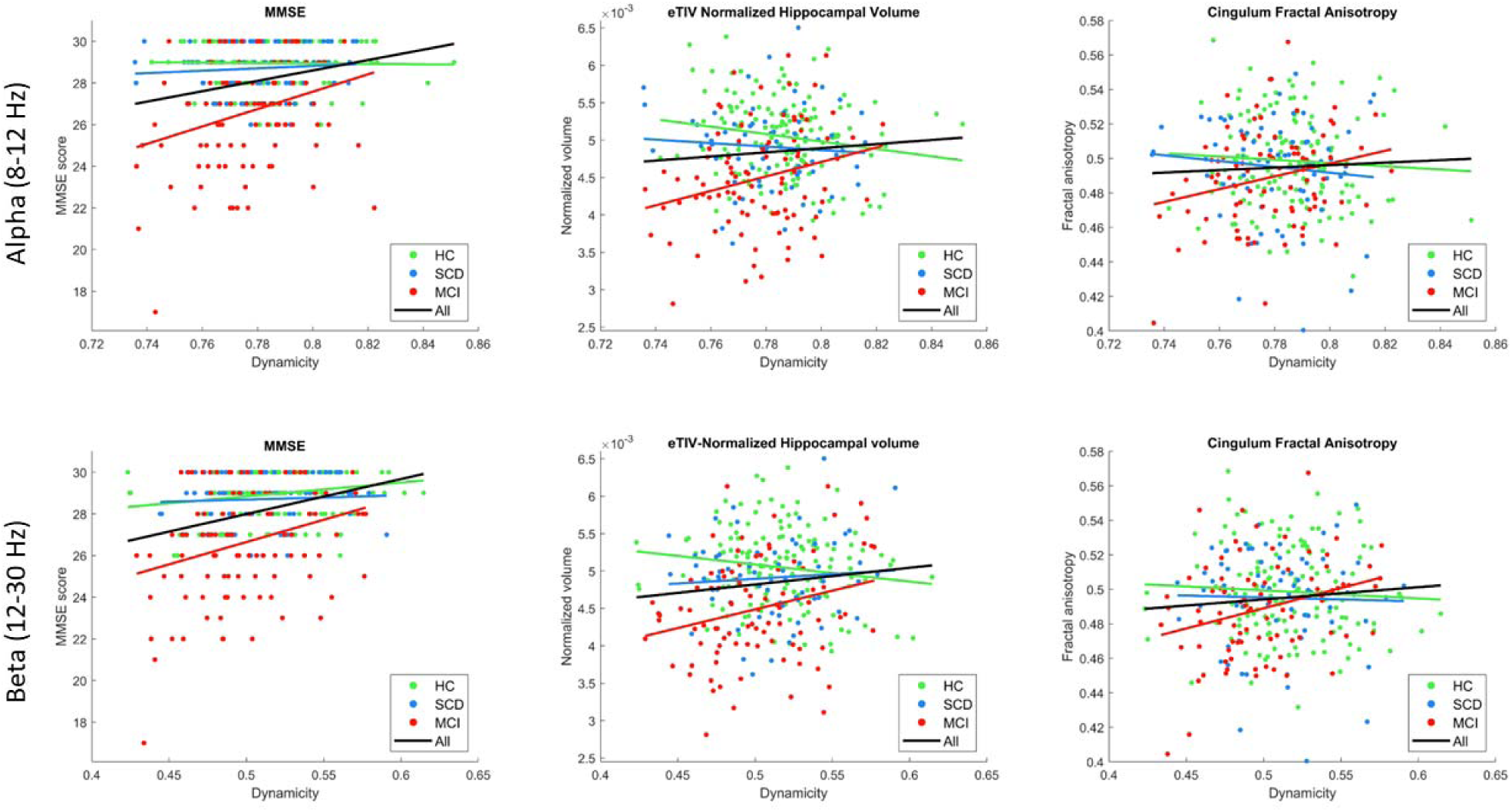
Correlations between whole-brain dynamicity and MMSE, eTIV-normalized hippocampal volume and the left Cingulum fractal anisotropy. Green corresponds to the HC group, blue to the SCD group, and red to the MCI group. The black line represents the correlation calculated when considering all subjects at the same time.

Interestingly, the MCI group showed multiple significant positive correlations, including those between dynamicity and MMSE scores, normalized hippocampus, and total grey matter volumes, and both left and right ILF tract FA. While the cingulum tracts FA did not maintain significance after FDR, both right and left tracts show a possible trend for a positive correlation with dynamicity.

At the same time, when calculating correlations with all the dataset, significant direct correlations between dynamicity and MMSE score, delayed logic recall score, normalized hippocampus and total cortical grey matter volume, and left cingulum FA were found. Both *rhos* and *corrected p-values* for all correlations calculated are shown in Table 2, where correlations found significant after FDR are marked in bold and with an asterisk.

**Table 2.** Correlation results between whole brain dynamicity and neuropsychological, anatomic and diffusion tensor imaging (DTI) parameters. Correlations were calculated for each group individually and then using all subjects in the sample (as indicated by the “All” column. Significant corrected p-values after FDR correction (q = 0.05) are bolded and marked with an asterisk. Column colours correspond to those assigned to each group in Figure 4.

| Alpha |  |  |  |  |  |  |  |  |
| --- | --- | --- | --- | --- | --- | --- | --- | --- |
|  | HC |  | SCD |  | MCI |  | All |  |
|  | R | P | R | P | R | P | R | P |
| MMSE | 0.0530 | 0.6084 | 0.0868 | 0.6446 | **0.2664** | **0.0234\*** | **0.2398** | **7.43E-6\*** |
| Delayed Logic | 0.0247 | 0.7725 | 0.0333 | 0.7230 | 0.0065 | 0.4756 | **0.1592** | **0.0088\*** |
| Hippocampus | -0.0570 | 0.7947 | -0.0297 | 0.8370 | **0.2653** | **0.0234\*** | **0.1378** | **0.0175\*** |
| Total grey | -0.0213 | 0.7947 | 0.0871 | 0.6446 | **0.2196** | **0.0356\*** | **0.1452** | **0.0140\*** |
| Left Cingulum | -0.0896 | 0.8457 | -0.0409 | 0.8370 | 0.1746 | 0.0748 | 0.0425 | 0.2531 |
| Right Cingulum | -0.0409 | 0.7947 | -0.1058 | 0.8545 | 0.1818 | 0.0712 | 0.0394 | 0.2537 |
| Left Inferior | 0.0880 | 0.4844 | -0.0708 | 0.8370 | **0.2047** | **0.0494\*** | 0.1038 | 0.0583 |
| Right Inferior | 0.1043 | 0.4844 | 0.1039 | 0.6446 | **0.2296** | **0.0356\*** | **0.1317** | **0.0233\*** |

| Beta |  |  |  |  |  |  |  |  |
| --- | --- | --- | --- | --- | --- | --- | --- | --- |
|  | HC |  | SCD |  | MCI |  | All |  |
|  | R | P | R | P | R | P | R | P |
| MMSE | **0.2747** | **0.0078\*** | 0.0147 | 0.7230 | **0.2550** | **0.0234\*** | **0.2787** | **4.55E-06\*** |
| Delayed Logic | 0.1129 | 0.4844 | 0.0497 | 0.7230 | 0.0296 | 0.4164 | **0.1939** | **0.0018\*** |
| Hippocampus | -0.0487 | 0.7947 | 0.1093 | 0.6446 | **0.2457** | **0.0313\*** | **0.1495** | **0.0132\*** |
| Total grey | -0.0514 | 0.7947 | 0.1704 | 0.6446 | **0.3043** | **0.0234\*** | **0.1737** | **0.0054\*** |
| Left Cingulum | -0.0281 | 0.7947 | 0.1388 | 0.6446 | **0.2307** | **0.0356\*** | **0.1232** | **0.0301\*** |
| Right Cingulum | -0.0429 | 0.7947 | -0.0777 | 0.8370 | **0.2246** | **0.0356\*** | 0.0574 | 0.1910 |
| Left Inferior | 0.0808 | 0.4844 | -0.1519 | 0.8882 | 0.1193 | 0.1722 | 0.0832 | 0.0994 |
| Right Inferior | 0.0801 | 0.4844 | 0.0280 | 0.7230 | 0.0972 | 0.2165 | 0.0887 | 0.0901 |

## Discussion

In this pioneering study, we aimed to evaluate the evolution of dFC measured with electrophysiology over the early AD spectrum. To this end, we performed the correlation of connectivity matrices over time, both globally and at source-level, a metric simple to calculate and interpret, and that only imposes assumptions regarding the size of the sliding window used for the calculations. Finally, we studied the relationship between this metric and the cognitive and anatomical hallmarks of AD. Our findings suggest that AD progression involves a reduction in dFC, mainly in the alpha and beta bands, which is especially pronounced over orbital and temporal areas and in regions involving the DMN. In addition, the decrease in dFC is related to cognitive decline and anatomical deterioration, both when considering data from all groups and, particularly, in the MCI group. These results shed light towards the understanding of brain dynamics along the AD continuum.

Whole-brain dFC analyses displayed a reduction of dynamicity over the continuum of AD, with differences of medium effect sizes between HCs and MCIs in the alpha (*Cohen’s d* = 0.474) and beta (*Cohen’s d* = 0.506) frequency bands, as shown in Figure 2. The SCD group did not show significant differences in their whole-brain dFC in any frequency band when compared to HCs or MCIs, proving to be an intermediate state between both conditions. This dFC decreases evidence that AD progression involves a reduction in connectivity profiles over time or, in other words, an increase in the consistency of connectivity patterns. These findings are aligned with previous literature on dFC, where a shrinkage in the dynamic repertoire of connectivity is observed through electrophysiological measurements^14,23^ and fMRI-based metrics.^16,17,20^ Additionally, studies focused on brain activity entropy, which depict the quantity of information on individual nodes of activity, find a decrease in signal complexity with the advance of the pathology,^24,49,50^ as well as in consciousness disorders.^51^ Remarkably, significant decreases in dynamicity between HC and MCI groups were observed specifically in the alpha and beta bands, agreeing with results in Núñez *et al*.^14^ and Li *et al*.,^23^ even if they report differences between HCs and AD participants, while we found those earlier in the AD continuum, between the HC and MCI groups. On a related note, recent studies have observed a dilution of FC topologies in MCI and AD participants, with decreasing modularity,^22^ and a blurring of frequency-dependent connectivity structure.^52^ This increase in frequency and spatial network homogeneity is thought to underlie the loss of specialized and integrated networks observed in these pathologic states and, in line with our results, could contribute to the decrease in temporal variability observed in the present study.

What physiological alterations may explain these changes in dFC? Previous literature has already discussed the reduction of the dynamic repertoire of connectivity along the spectrum of AD,^16^ linking it to possible alterations in white matter integrity. AD is associated with myelin aberrations both in animal models^53^ and in humans^54^, which may occur as early as 20 years before the symptoms’ onset, and has been also been connected to β-amyloid deposition.^55^ At the same time, several studies have drawn a connection between myelin abnormalities and brain connectivity, highlighting the importance of tract myelination, transmission speed, and the resulting coupling and synchrony patterns observed in the brain.^56,57^ More specifically, it has been observed that the relations between structural and FC are spatial and frequency-dependent^58,59^ and varies with age.^60^ Importantly, multiple works, such as Karahan and colleagues,^58^ have established a connection between the frequency bands affected in this study and myelination distribution and white matter properties. In their work, they showed negative correlations between beta sFC inter-subject variability (ISV) and myelination, with alpha sFC-ISV failing to maintain significance due to statistical corrections, and significant negative correlations between alpha and beta sFC-ISV and the hindrance of white matter pathways. Additionally, the temporal precision needed for the oscillatory coupling and phase-locking increases with oscillation frequency, leading to enhanced susceptibility to myelin degradation for higher frequency bands.^59^ Our results are aligned with all their conclusions, as we found increased effect sizes of differences over the whole-brain and source-level dFC and more widespread differences between MCI and HC groups over the beta band than in the alpha band, while not finding any result in the theta band. These events support the theory of increased vulnerability of brain coupling at higher frequencies. We also discovered significant negative correlations between whole-brain beta and alpha dFC and FA in multiple tracts, indicating a strong and important dependence between white matter degradation and network integrity in AD.

The relevance of the regions and brain networks found using seed-based dFC (including orbitofrontal and temporal areas for both alpha and beta bands, and also precuneus, IFG, anterior and posterior cingulate gyrus, and superior medial frontal gyrus for the latter) has been highlighted in former research on brain changes throughout the AD continuum, such as functional alterations,^2,6,8,9,23^ volumetric atrophy observed with MRI,^61,62^ and amyloid neuropathological disturbances measured with positron emission tomography.^63^ Interestingly, the distribution of the dFC changes found between MCI and HC correspond to multiple results in Karahan et al.^58^ Brain areas with decreased hindrance of white matter pathways display a similar distribution as the significant cluster found in the alpha band, and the location of the areas with the lowest cortical R1, e.g., with the lowest myelination, is alike the cluster found in the beta band. These areas are characterized by a lower neuronal density, large dendritic arborization, more spine density, more synapses, and higher aerobic glycolysis, which may be a neurobiological feature of the association cortices, enabling adaptable and plastic neural circuitry. These attributes might, in turn, lead to more complex FC behaviors, highlighting the importance of loss of dynamic activity in these areas along the AD continuum.

Finally, the potential of the proposed dFC marker is indicated by its significant correlations in alpha and beta with the cognitive and anatomical parameters. It is worth highlighting that those trends were found both when using the whole dataset and for the MCI group only. While correlations found in the whole dataset showcase the value of dynamicity as a possible biomarker for AD progression, it includes groups with significantly different cognitive and anatomical scores. On the other hand, MCI group correlations imply that lower dFC is related to the severity of the signs these participants show, as well as showing the link between dFC and anatomical damage due to the advance of the disease. Surprisingly, we can find studies describing significant relationships between sFC and DTI measures,^64–66^ and at the same time alternative sFC-based investigations that do not find a relationship between these two markers.^67–69^ Based on a relatively large sample, the correlations found in this study between dFC and white matter degradation in the pre-dementia phase of the AD continuum contrast with those reported using sFC and motivate further investigation to unveil the potential sensitivity of dFC to early structural changes in the brain.

Nevertheless, some limitations need to be noted. Firstly, the use of a sliding-window technique does not come without disadvantages. The size of the window used in this study (4 seconds) has been used in previous relevant studies determining relevant FC features in AD for the frequency bands investigated here,^6,8–10,70^ but it was not large enough to capture sufficient oscillations in the delta band, and not small enough to capture dynamic patterns in the gamma band. Future studies will address this limitation for the study of the latter frequency band, especially relevant for AD, and according to our discussion particularly vulnerable to myelin damage. While based on a large dataset, this study did not include any participant with AD, necessary to obtain a complete assessment of the entire AD spectrum. Future studies should investigate the relation of dFC with white matter integrity, sFC, and neural excitation/inhibition.

## Conclusion

In conclusion, our study offers compelling evidence that dFC declines across the AD spectrum, particularly within the alpha and beta frequency bands, emphasizing its potential as a biomarker for early AD detection and progression monitoring. This decline in dFC, especially pronounced in critical brain regions involved in the default mode network, correlates with cognitive deterioration and structural brain changes, further indicating the relevance of dFC in AD, and underpinning the interconnected nature of functional, cognitive, and anatomical alterations in this disorder.

## Acknowledgements

We would like to thank all participants that were included in this study for their selfless contribution to science, making this work possible.

## Data availability

The data and the algorithms that support the findings of this study are available from the corresponding author, upon reasonable request.

## IRB statement

The “Hospital Clínico San Carlos” Ethics Committee approved this study, and the procedure was performed following internationally accepted guidelines and regulations. Moreover, signed consent was obtained from all participants.

