## Supplementary Table 1 for "Dynamics of Brain Connectivity across the Alzheimer’s Disease Spectrum: a magnetoencephalography study"

**Table S1. Sample demographics of the stable MCI (sMCI) and progressive MCI (pMCI) participants.** FDR corrected p-values for the comparisons are shown in the rightmost column.

| | **sMCI** | **pMCI** | **p-value** |
|---|---|---|---|
| N | 64 | 42 | |
| Sex | 20M/44F | 18M/26F | 0.1489 |
| Age | 73.5±5.3 | 75.7±4.8 | 0.5924 |
| MMSE | 27.0±2.4 | 25.9±2.9 | 0.1988 |
| Delayed recall test | 6.8±7.1 | 2.3±4.3 | 0.1489 |
| Total grey matter volume (eTIV-normalized) | 0.290±0.020 | 0.270±0.021 | 0.0700 |
| Hippocampus volume (eTIV-normalized) | 0.0047±0.0006 | 0.0041±0.0006 | **2.68E-04*** |
| Left Cingulum tract FA | 0.522±0.036 | 0.515±0.030 | 0.3838 |
| Right Cingulum tract FA | 0.491±0.028 | 0.486± 0.028 | 0.1489 |
| Left Inferior Logitudinal tract FA | 0.491±0.025 | 0.470±0.090 | 0.1489 |
| Right Inferior Longitudinal tract FA | 0.496±0.028 | 0.471±0.091 | 0.1489 |
