## Supplementary Table 2 for "Dynamics of Brain Connectivity across the Alzheimer’s Disease Spectrum: a magnetoencephalography study"

**Table S2. Complete statistical report on the whole-brain dynamicity between-group linear regression.** Significant FDR corrected (q = 0.05) p-values are bolded and have an asterisk. Significant effects of the "Group" variable were found in the alpha and beta bands when comparing HC and MCI groups, with their corrected p-values highlighted in yellow. SE: standard error; HC: Healthy control, SCD: Subjective cognitive decline, MCI: Mild cognitive impairment.

| | | HC vs SCD | | | | HC vs MCI | | | | SCD vs MCI | | | |
|---|---|---|---|---|---|---|---|---|---|---|---|---|---|
| | | **β-value** | **SE** | ***t* -Stat** | **p-value** | **β-value** | **SE** | ***t* -Stat** | **p-value** | **β-value** | **SE** | ***t* -Stat** | **p-value** |
| **Theta** | Intercept | 0.721303 | 0.021914 | 32.91510 | 3.34E-84* | 0.72369 | 0.019671 | 36.78913 | 2.77E-100* | 0.761485 | 0.021495 | 35.42653 | 1.97E-80* |
| | Group | -0.00294 | 0.002822 | -1.04144 | 0.3586 | -0.00325 | 0.002619 | -1.24378 | 0.26662 | 0.001748 | 0.003175 | 0.55065 | 0.65541 |
| | Age | 0.000655 | 0.000302 | 2.16831 | 0.05358 | 0.00060 | 0.000273 | 2.21726 | 0.049550* | 3.52E-05 | 0.000294 | 0.11972 | 0.93069 |
| | Sex | -0.0098 | 0.002853 | -3.43442 | 0.00214* | -0.00794 | 0.002558 | -3.10718 | 0.00543* | -0.00847 | 0.003247 | -2.60984 | 0.02089* |
| **Alpha** | Intercept | 0.764453 | 0.019991 | 38.24012 | 9.38E-96* | 0.76037 | 0.017675 | 43.0197 | 2.37E-114* | 0.785415 | 0.01949 | 40.29878 | 6.78E-89* |
| | Group | -0.00372 | 0.002575 | -1.44426 | 0.20019 | -0.00989 | 0.002353 | -4.20261 | **1.33E-04*** | -0.00488 | 0.002879 | -1.69468 | 0.13239 |
| | Age | 0.000366 | 0.000276 | 1.32796 | 0.23866 | 0.00042 | 0.000245 | 1.729969 | 0.12735 | 2.01E-05 | 0.000267 | 0.07522 | 0.94012 |
| | Sex | -0.00734 | 0.002603 | -2.82163 | 0.01256* | -0.00750 | 0.002298 | -3.26365 | 0.00348* | -0.00741 | 0.002944 | -2.51699 | 0.02549* |
| **Beta** | Intercept | 0.511855 | 0.037904 | 13.50388 | 9.29E-30* | 0.50679 | 0.034789 | 14.56774 | 2.66E-34* | 0.581195 | 0.037565 | 15.4717 | 8.32E-34* |
| | Group | -0.00413 | 0.004882 | -0.84550 | 0.46311 | -0.01865 | 0.004632 | -4.02662 | **0.00025*** | -0.01098 | 0.005549 | -1.97834 | 0.07744 |
| | Age | 0.000171 | 0.000523 | 0.32809 | 0.78688 | 0.00024 | 0.000483 | 0.500797 | 0.67305 | -0.00086 | 0.000514 | -1.66853 | 0.13433 |
| | Sex | -0.01181 | 0.004935 | -2.39289 | 0.03333* | -0.01176 | 0.004524 | -2.60091 | 0.02089* | -0.01204 | 0.005674 | -2.12128 | 0.05780 |
