## Supplementary Table 3 for "Dynamics of Brain Connectivity across the Alzheimer’s Disease Spectrum: a magnetoencephalography study"

**Table S3. Complete statistical report from the stable MCI (sMCI) and progressive MCI (pMCI) linear regressions.** No grouping variable had a significant effect after FDR correction (q = 0.05).

| | | sMCI vs pMCI | | | |
|---|---|---|---|---|---|
| | | **β-value** | **SE** | **t -Stat** | **p-value** |
| **Theta** | Intercept | 0.721303 | 0.021914 | 32.91511 | 3.20-46* |
| | Group | -0.00294 | 0.002822 | -1.04145 | 0.9607 |
| | Age | 0.000655 | 0.000302 | 2.168316 | 0.8888 |
| | Sex | -0.0098 | 0.002853 | -3.43443 | 0.2729 |
| **Alpha** | Intercept | 0.764453 | 0.019991 | 38.24012 | 5.60E-51* |
| | Group | -0.00372 | 0.002575 | -1.44427 | 0.4833 |
| | Age | 0.000366 | 0.000276 | 1.32797 | 0.6584 |
| | Sex | -0.00734 | 0.002603 | -2.82164 | 0.1050 |
| **Beta** | Intercept | 0.511855 | 0.037904 | 13.50389 | 1.80E-17* |
| | Group | -0.00413 | 0.004882 | -0.84551 | 0.6584 |
| | Age | 0.000171 | 0.000523 | 0.328092 | 0.6584 |
| | Sex | -0.01181 | 0.004935 | -2.3929 | 0.2729 |
