## Supplementary Table 4 for "Dynamics of Brain Connectivity across the Alzheimer’s Disease Spectrum: a magnetoencephalography study"

**Table S4. List of the regions of interest (ROI) of the Automated Anatomical Labeling (AAL) atlas contained in the significant cluster found between the HC and MCI groups in the alpha band.** Only the ROIs whose volume overlapped at least 10% with the cluster were included in the list. Green corresponds to frontal lobe areas, blue to temporal lobe areas and magenta to limbic lobe areas.

| Area Name | % of ROI contained in the cluster | % of cluster |
|---|---|---|
| Left Middle Frontal gyrus, Orbital | 100.00% | 8.97% |
| Left Inferior Temporal gyrus | 29.17% | 8.97% |
| Right Middle Frontal gyrus, Orbital | 85.71% | 7.69% |
| Left Superior Frontal gyrus, Medial Orbital | 100.00% | 7.69% |
| Left Middle temporal gyrus | 13.64% | 7.69% |
| Left Inferior Frontal gyrus, Orbital | 41.67% | 6.41% |
| Right Superior Frontal gyrus, Medial Orbital | 100.00% | 6.41% |
| Left Temporal pole, Superior Temporal gyrus | 50.00% | 6.41% |
| Left Temporal pole, Middle temporal gyrus | 71.43% | 6.41% |
| Left Gyrus Rectus | 50.00% | 5.13% |
| Left Fusiform gyrus | 26.67% | 5.13% |
| Left Superior Frontal gyrus | 11.11% | 3.85% |
| Left Superior Frontal gyrus, Orbital | 66.67% | 2.56% |
| Right Superior Frontal gyrus, Orbital | 66.67% | 2.56% |
| Right Gyrus Rectus | 50.00% | 2.56% |
| Left Cingulate gyrus, Anterior part | 10.53% | 2.56% |
| Left Parahippocampus | 25.00% | 2.56% |
| Left Hippocampus | 20.00% | 1.28% |
