## Supplementary Table 5 for "Dynamics of Brain Connectivity across the Alzheimer’s Disease Spectrum: a magnetoencephalography study"

**Table S5. List of the regions of interest (ROI) of the Automated Anatomical Labeling (AAL) atlas contained in the significant cluster found between the HC and MCI groups in the beta band.** Only the ROIs whose volume overlapped at least 10% with the cluster were included in the list. Blue corresponds to temporal lobe areas, yellow to parietal lobe areas, green to frontal lobe areas, magenta to limbic lobe areas, and red to occipital lobe areas.

| Area Name | % of ROI contained in the cluster | % of cluster |
|---|---|---|
| Right Superior Temporal gyrus | 81.48% | 9.40% |
| Left Middle temporal gyrus | 34.09% | 6.41% |
| Left Precuneus | 50.00% | 5.98% |
| Left Superior Temporal gyrus | 70.00% | 5.98% |
| Right Postcentral gyrus | 39.39% | 5.56% |
| Left Inferior Temporal gyrus | 50.00% | 5.13% |
| Left Inferior Frontal gyrus, Orbital | 66.67% | 3.42% |
| Right Middle temporal gyrus | 21.62% | 3.42% |
| Left Cingulate gyrus, Middle part | 43.75% | 2.99% |
| Left Temporal pole, Superior Temporal gyrus | 70.00% | 2.99% |
| Left Middle Frontal gyrus, Orbital | 85.71% | 2.56% |
| Right Rolandic operculum | 54.55% | 2.56% |
| Left Insula | 42.86% | 2.56% |
| Left Calcarine fissure and surrounding cortex | 30.00% | 2.56% |
| Right Supramarginal gyrus | 60.00% | 2.56% |
| Right Paracentral lobule | 75.00% | 2.56% |
| Left Temporal pole, Middle temporal gyrus | 85.71% | 2.56% |
| Left Superior Frontal gyrus, Medial Orbital | 83.33% | 2.14% |
| Right Insula | 35.71% | 2.14% |
| Left Cingulate gyrus, Anterior part | 26.32% | 2.14% |
| Right Superior Parietal gyrus | 27.78% | 2.14% |
| Right Precuneus | 23.81% | 2.14% |
| Left Gyrus Rectus | 50.00% | 1.71% |
| Left Hippocampus | 80.00% | 1.71% |
| Left Fusiform gyrus | 26.67% | 1.71% |
| Right Inferior Parietal gyrus | 36.36% | 1.71% |
| Left Supplementary Motor area | 12.50% | 1.28% |
| Left Cingulate gyrus, Posterior part | 60.00% | 1.28% |
| Left Superior Frontal gyrus, Orbital | 66.67% | 0.85% |
| Left Rolandic operculum | 40.00% | 0.85% |
| Left Olfactory cortex | 100.00% | 0.85% |
| Right Superior Frontal gyrus, Medial Orbital | 40.00% | 0.85% |
| Left Parahippocampus | 25.00% | 0.85% |
| Left Cuneus | 18.18% | 0.85% |
| Right Heschl's gyrus | 100.00% | 0.85% |
| Right Olfactory cortex | 25.00% | 0.43% |
| Right Gyrus Rectus | 25.00% | 0.43% |
| Left Paracentral lobule | 11.11% | 0.43% |
| Left Heschls gyrus | 33.33% | 0.43% |
| Right Temporal pole, Superior Temporal gyrus | 12.50% | 0.43% |
