## Supplementary Figure 1 for "Dynamics of Brain Connectivity across the Alzheimer’s Disease Spectrum: a magnetoencephalography study"

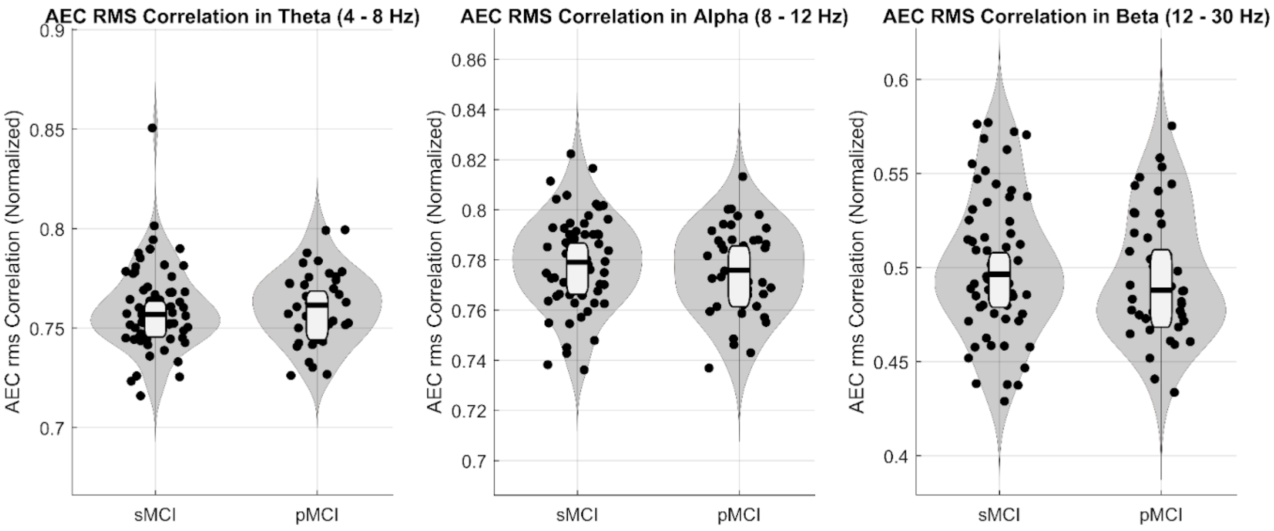


**Figure S1. Violin plots depicting whole-brain dynamicity in the stable MCI (sMCI) and progressive MCI (pMCI) groups.** No significant differences were found.
